# CREB mediates the *C. elegans* dauer polyphenism through direct and cell-autonomous regulation of TGF-β expression

**DOI:** 10.1101/2020.08.14.250498

**Authors:** Jisoo Park, Hyekyung Oh, Do-Young Kim, YongJin Cheon, Yeon-Ji Park, Scott J. Neal, Abdul Rouf Dar, Rebecca A. Butcher, Piali Sengupta, Daewon Kim, Kyuhyung Kim

## Abstract

Animals can adapt to dynamic environmental conditions by modulating their developmental programs. Understanding the genetic architecture and molecular mechanisms underlying developmental plasticity in response to changing environments is an important and emerging area of research. Here, we show a novel role of cAMP response element binding protein (CREB)-encoding *crh-1* gene in developmental polyphenism of *C. elegans*. Under conditions that promote normal development in wild-type animals, *crh-1* mutants inappropriately form transient pre-dauer (L2d) larva and express the L2d marker gene. L2d formation in *crh-1* mutants is specifically induced by the ascaroside pheromone ascr#5 (asc-ωC3; C3), and *crh-1* functions autonomously in the ascr#5-sensing ASI neurons to inhibit L2d formation. Moreover, we find that CRH-1 directly binds upstream of the *daf-7* TGF-β locus and promotes its expression in the ASI neurons. Taken together, these results provide new insight into how animals alter their developmental programs in response to environmental changes.

## Introduction

Animals exhibit phenotypic traits whose expression can be influenced by environmental signals, including temperature, food availability, and crowding. These signals are sensed, processed, and integrated by the nervous system, and govern appropriate developmental and/or behavioral phenotypes through effecting local and systemic changes in gene expression and/or endocrine signals. For example, population density and food supply determine wing development (winged vs. wingless) of the pea aphid (*A. pisum*) via ecdysone signaling (Vellichirammal, Gupta et al., 2017). This extreme form of phenotypic plasticity is referred to as polyphenism -the development of alternate and distinct phenotypes in animals of the same genotype - and is found across taxa (Fusco & Minelli, 2010, Projecto-Garcia, Biddle et al., 2017). Although the ability of animals to modulate their developmental programs in response to changing environmental conditions is crucial for their survival, the molecular mechanisms underlying such developmental plasticity are not yet fully understood.

The nematode *Caenorhabditis elegans* exhibits polyphenic development that is regulated by environmental conditions. Newly hatched L1/L2 larvae of *C. elegans* sense and integrate environmental signals, including food availability, temperature, and the abundance of pheromones (a population density indicator) to determine whether to undergo normal reproductive development into the L3 larval stage, or arrest at an alternate L3 stage, known as dauer. Dauer larvae are developmentally arrested, non-aging, stress-resistant, and can re-enter normal reproductive development if conditions improve (Cassada & Russell, 1975, Golden & Riddle, 1984) (Fig 1A). Under less favorable conditions, L1 larvae may transiently develop as L2d larvae, an obligate precursor to dauer commitment, but which still retain the capacity to resume reproductive development (Golden & Riddle, 1984) (Fig 1A). Thus, worms assess and weigh multiple factors at multiple time points along the dauer trajectory in making the dauer decision. However, it is not clear how worms evaluate external and internal signals at specific developmental time windows and execute dauer formation. Specifically, the signals and molecules that promote the L1-to-L2d developmental transition are also not well understood.

**Figure 1.**
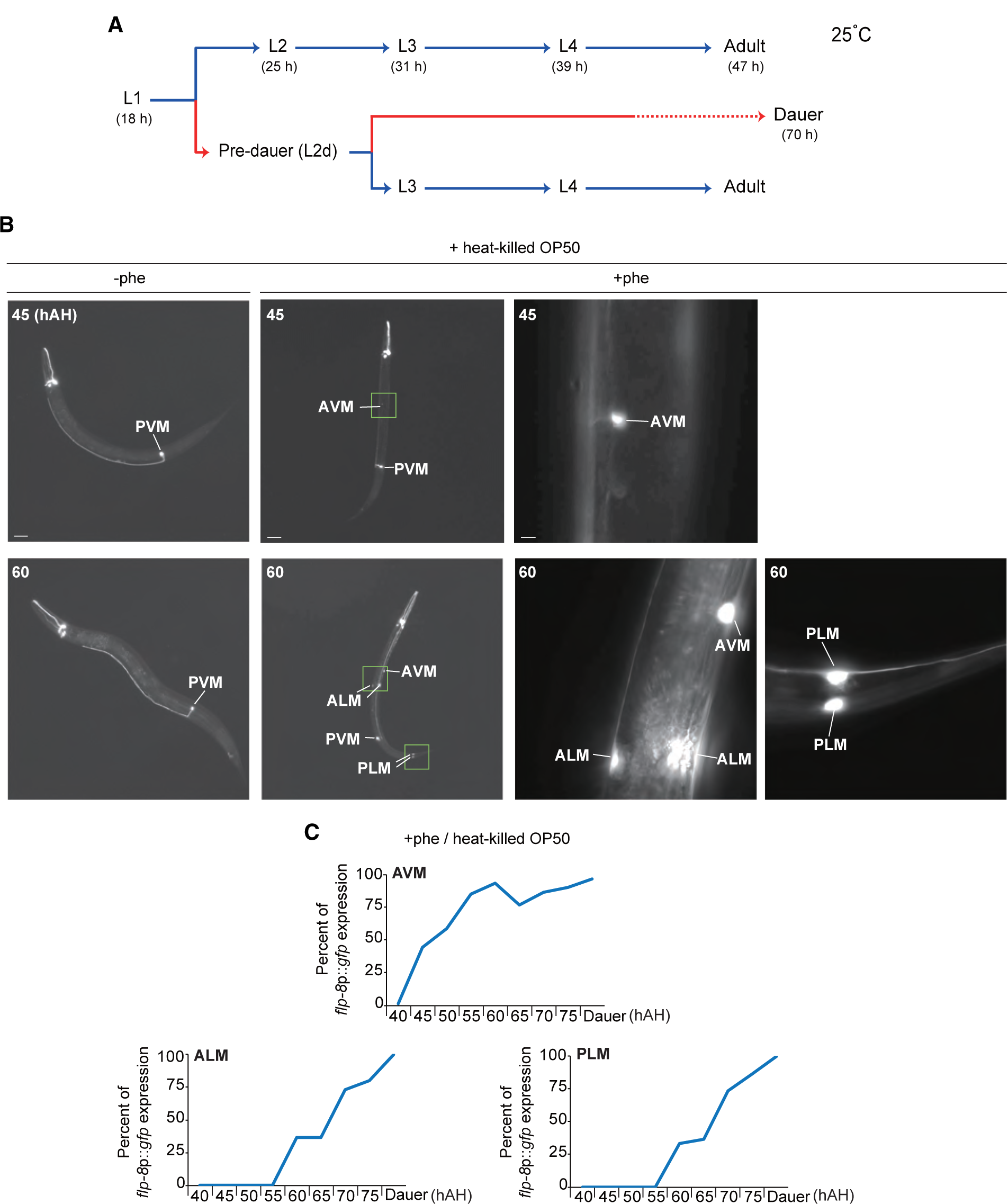
The *flp-8* expression is turned on in a subset of touch receptor neurons of L2d larvae. **A**. *C. elegans* life cycle at 25°C. In favorable conditions, newly hatched L1/L2 larvae undergo normal reproductive development into L3 larvae. Under unfavorable conditions, L1 larvae arrest as dauer larvae via the L2d pre-dauer stage. Under less favorable conditions, L1 larvae transiently become L2d larvae, which can resume reproductive growth. Numbers show the approximate duration in hours of development of each stage. **B**. Representative images of wild-type animals expressing of *flp-8*p::*gfp* in a subset of touch receptor neurons in the presence of heat-killed OP50 food without (left column) or with (right three columns) ascr#5 pheromone. The boxed regions of the second column are shown on the right two columns at higher magnification. hAH, hour after hatching. Scale bars: 50 µm (left two columns), 5 µm (right two columns). **C**. Percent of *flp-8*p::*gfp* expression in the AVM, ALM and PLM neurons by wild-type animals when grown in the presence of heat-killed OP50 food and ascr#5 pheromone. n ≥ 30 for each.

As a dauer-inducing cue, *C. elegans* secretes pheromones, which are a complex cocktail of small molecules called ascarosides (Butcher, Fujita et al., 2007, Butcher, Ragains et al., 2008, Jeong, Jung et al., 2005, Srinivasan, Kaplan et al., 2008). Moreover, worms produce and secrete distinct concentrations and combinations of ascarosides depending upon developmental stages and environmental conditions (Butcher et al., 2008, Kaplan, Srinivasan et al., 2011, von Reuss, Bose et al., 2012). Although the potency and property of pheromone components in inducing dauer formation have been described, little is known about the contribution of each component to specific time windows along the dauer trajectory.

The molecular and genetic mechanisms underlying the decision to enter the dauer stage have been extensively studied, and have established two neuroendocrine *daf-7*/TGF-β and *daf-2*/insulin signaling pathways as critical mediators of this binary fate decision (Kimura, Tissenbaum et al., 1997, Ren, Lim et al., 1996, Schackwitz, Inoue et al., 1996). Expression of *daf-7* in the ASI sensory neuron type is regulated by environmental signals, including temperature, food availability, pheromone, and infection (Kim, Sato et al., 2009, Meisel, Panda et al., 2014, Ren et al., 1996, Schackwitz et al., 1996). However, the mechanisms that regulate *daf-7* expression as a function of environmental conditions have not been fully described.

*crh-1*-encoded CREB (cAMP response element binding protein) plays critical roles in *C. elegans* metabolism, sensory responses, lifespan, and learning and memory (Chen, Chen et al., 2016). Similar to mammalian CREB, CRH-1 directly regulates target gene expression via the evolutionarily conserved cAMP-responsive element (CRE) and represents a nutritive or metabolic indicator (Altarejos & Montminy, 2011). However, a role for CREB in developmental plasticity, such as the decision to enter the dauer stage in *C. elegans*, has not been explored.

Here, we show that CRH-1 plays a role in the decision to enter the L2d stage. In rich food conditions, pheromone is generally insufficient to induce L2d/dauer formation in wild-type animals. However, *crh-1* mutants exhibit transient L2d formation and express the L2d marker gene *flp-8* under such conditions. L2d formation of *crh-1* mutants is induced by ascr#5 (asc-ωC3; C3), but not by ascr#2 (asc-C6MK; C6) or ascr#3 (asc-ΔC9; C9), and expression of CRH-1 in the ascr#5-sensing ASI neurons is sufficient to rescue inappropriate L2d formation in *crh-1* mutants. We find that CRH-1 acts cell-autonomously in the ASI sensory neurons to regulate *daf-7* TGF-β expression and that ASI-specific overexpression of *daf-7* is sufficient to block L2d formation in *crh-1* mutants. We also identify a functional CRE in *daf-7* regulatory sequences and show that it is specifically bound by CRH-1, confirming that CRH-1 regulates *daf-7* expression in ASI. Together, these results reveal that CREB modulates developmental plasticity by repressing L2d entry in the presence of rich food sources and abundant ascr#5, via direct regulation of *daf-7* TGF-β expression in the ascr#5-sensing ASI neurons.

## Results

### A *flp-8* FMRFamide-like neuropeptide gene is expressed in a subset of touch receptor neurons in pre-dauer larvae

To better understand the mechanisms underlying L2d larvae formation, we first screened for a marker that specifically identifies the L2d developmental stage. The *flp-8* FMRFamide-like gene was previously shown to be differentially expressed in a subset of touch receptor neurons (AVM, ALM, PLM; referred to as AAP-TRN henceforth) in dauer larvae in addition to expression in the AUA, URX and PVM neurons throughout larval development (Kim & Li, 2004). We monitored the onset and maintenance of *flp-8*p::*gfp* reporter gene expression in the AAP-TRN of animals grown under non-dauer-inducing conditions and dauer-inducing conditions with limited heat-killed OP50 food and with and without the ascr#5 pheromone, respectively (Butcher, 2017). Consistent with the previous study (Kim & Li, 2004), we detected the expression of *flp-8*p::*gfp* in AVM only in the dauer-inducing conditions. Expression in AVM was first detected 40 h after hatching (hAH) with more prominent expression evident at 60 hAH; 60 hAH is approximately 20 h prior to dauer entry/commitment under these conditions (Fig 1B-C). Animals expressing *flp-8*p::*gfp* in the AVM neurons at 60 hAH were morphologically similar to dauer larvae but did not survive under 1% sodium dodecyl sulfate (SDS) (Cassada & Russell, 1975) treatment (n ≥ 50), implying that *flp-8* expression in the AVM neurons begins at the L2d stage, before the completion of dauer cuticular remodeling that confers resistance to this detergent. We noted the onset of *flp-8* expression in the ALM and PLM neurons at 55 hAH, five hour prior to dauer entry (Fig 1B-C). All dauer larvae expressed *flp-8*p::*gfp* in the AAP-TRN (Fig 1B-C). Taken together, these results suggest that the onset of *flp-8*p::*gfp* expression in the AVM neurons and possibly the ALM and PLM neurons can be used as a marker for the L2d stage.

### *crh-1* CREB mutants express an *flp-8* L2d marker gene in non-dauer-inducing conditions

We next screened for mutants with altered *flp-8* expression. In a specific non-dauer-inducing condition with abundant live OP50 food and high concentrations of ascr#5, *crh-1* mutant larvae inappropriately expressed the *flp-8* reporter gene in the AAP-TRN (Fig 2A-B). However, in dauer-inducing conditions with limited heat-killed OP50 food and high concentrations of ascr#5 pheromone, the expression of *flp-8* in *crh-1* mutant larvae in AAP-TRN was similar to that of wild-type, although a slightly smaller percentage of *crh-1* mutants expressed GFP in the AAP-TRN as compared to wild-type (Fig EV1). All assays were conducted in this non-dauer inducing condition with abundant live OP50 food unless noted otherwise. The onset of expression in these neurons in *crh-1* mutants under non dauer-inducing conditions was approximately 35 hAH, five hours earlier than in the dauer-inducing condition, likely due to their rapid growth from having live food available (Croll, Smith et al., 1977). *flp-8* expression was not observed in the AAP-TRN of wild-type or *crh-1* mutant larvae in the absence of pheromone (Fig EV2), indicating that the exposure to pheromone is necessary for induction of *flp-8* expression in the AAP-TRN. These results show that *crh-1* mutants inappropriately express a L2d marker gene in conditions that do not promote dauer formation in wild-type animals.

**Figure 2.**
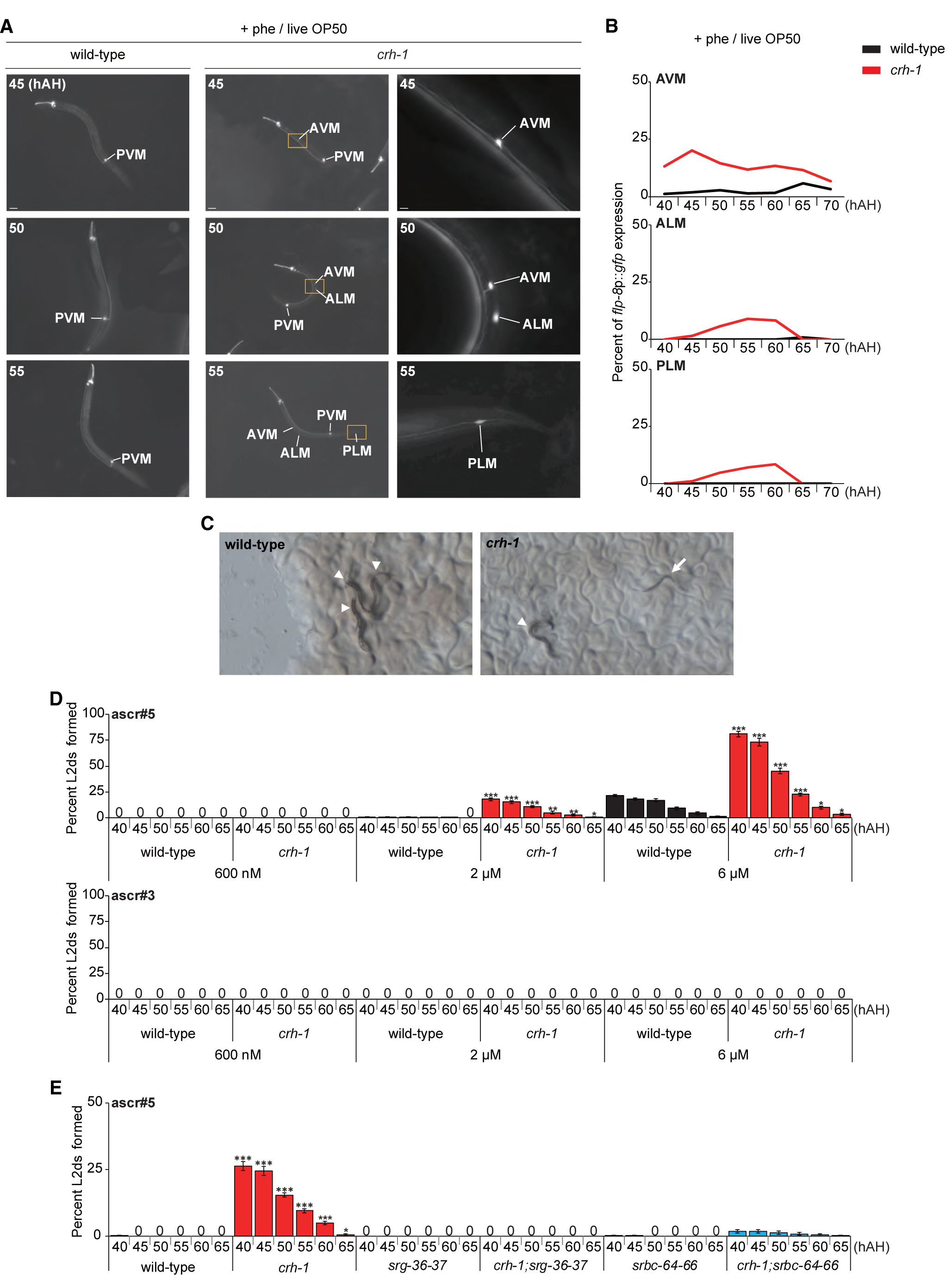
*crh-1* mutants inappropriately express an *flp-8* L2d marker gene and form transient pre-dauer L2d larvae in non-dauer inducing conditions. **A**. Representative images of wild-type (left column) or *crh-1* mutant (right two columns) animals expressing of *flp-8*p::*gfp* in a subset of touch receptor neurons in the presence of live OP50 food and ascr#5 pheromone. The boxed regions of the second column are shown on the right column at higher magnification. hAH, hour after hatching. Scale bars: 50 µm (left two columns), 5 µm (right column). **B**. Percent of *flp-8*p::*gfp* expression in the AVM, ALM and PLM neurons by wild-type or *crh-1* mutant animals when grown in the presence of live OP50 food and ascr#5 pheromone. n ≥ 30 each. **C**. Light microscopy images of wild-type (left) or *crh-1* mutant (right) animals in the presence of live OP50 and crude pheromone. Arrowhead or arrow indicates a L3 or L2d larva, respectively. **D**. Percent of L2d formed by wild-type or *crh-1* mutant animals when grown in the presence of live OP50 food and ascr#5 (top) or ascr#3 (bottom) pheromone. N ≥ 5 for each. Error bars indicate SEM. *\*, \*\**, and *\*\*\** indicate different from wild-type at p < 0.05, p < 0.01, and p < 0.001, respectively (student t-test). **E**. Percent of L2d formed by animals of the indicated genotypes when grown in the presence of live OP50 food and ascr#5 pheromone. N ≥ 8 for each. Error bars indicate SEM. *\** and *\*\*\** indicate different from wild-type at p < 0.05 and p < 0.001, respectively (one-way ANOVA with Bonferroni’s post hoc tests).

### *crh-1* mutants inappropriately form transient pre-dauer L2d larvae upon exposure to ascr#5

To better understand the *crh-1* mutant phenotype, we next examined the ability of *crh-1* mutants to enter the dauer stage in response to different concentrations of ascaroside pheromones, under standardized dauer-inducing conditions. We found that *crh-1* mutants exhibited normal or weakly decreased pheromone-mediated dauer formation (Fig EV3). However, in a non-dauer inducing condition with abundant live OP50 food and high concentrations of crude pheromone, *crh-1* mutants formed thin dauer-like larvae 40-60 hAH (Fig 2C, Fig EV4) consistent with inappropriate induction of the L2d stage. To assess whether these larvae were indeed dauers, we treated them with 1% SDS. None of the *crh-1* dauer-like larvae survived under 1% SDS treatment (n ≥ 50), indicating that these larvae have not fully remodeled their cuticle as expected for dauer larvae. We next characterized their morphology using scanning electron microscopy. The mouths of *crh-1* dauer-like larvae remained open as in L3 larvae (Fig EV5), suggesting, in combination with their other phenotypic attributes, that they are indeed L2d larvae. Moreover, by 65 hAH, these larvae had resumed reproductive development, consistent with transient passage through the L2d state and incongruent with dauer commitment and exit.

To determine whether the transient L2d formation of *crh-1* mutants is induced by specific pheromone components, we tested several ascaroside pheromones, including ascr#2, ascr#3, and ascr#5. All assays were conducted beginning 40 hAH unless noted otherwise. We found that *crh-1* mutants specifically formed transient L2d in the presence of micromolar concentrations of ascr#5, but not ascr#2 or ascr#3 (Fig 2D, Fig EV6). Moreover, *flp-8* was not expressed in the AAP-TRN of *crh-1* mutant larvae in response to ascr#2 and ascr#3 (Fig EV7).

ascr#5-mediated dauer formation has been shown to be primarily mediated by a pair of GTP-binding protein (G protein)-coupled receptors (GPCRs), SRG-36 and SRG-37 expressed specifically in ASI, and partly by another pair of GPCRs, SRBC-64 and SRBC-66 expressed in the ASK sensory neurons (Kim et al., 2009, McGrath, Xu et al., 2011). We found that the transient L2d formation of *crh-1* mutants was suppressed completely by mutations in both *srg-36* and *srg-37* and partially by mutations in both *srbc-64* and *srbc-66* (Fig 2E), supporting the specificity of ascr#5 in the transient L2d formation of *crh-1* mutants. Taken together, these results imply that *crh-1* mutants inappropriately and transiently enter into the L2d stage in response to ascr#5 prior to resuming reproductive development.

### *crh-1* acts in the ASI neurons to regulate ascr#5-mediated L2d formation

To identify the molecular mechanisms by which CRH-1 regulates ascr#5-mediated L2d formation, we identified the site of action of CRH-1. Previously, GFP driven by *crh-1* promoters had been shown to be expressed in a set of head neurons of adult *C. elegans* (Kimura, Corcoran et al., 2002). To further verify the expression patterns of *crh-1*, we generated transgenic animals expressing GFP under the control of sequences 2601 bp, 2411 bp, 1094 bp, or 737 bp (*crh-1*p*1*::*gfp* to *crh-1*p*4*::*gfp*) upstream of the translation start sequence of *crh-1* gene. We found that sequences 2601, 2411, and 1094 bp regulatory sequences drove expression specifically in three pairs of sensory neurons in the head amphid organs, including in the AWC, ASE, and AFD neurons, consistent with previous observations (Chen et al., 2016). However, we also routinely detected weak GFP expression in the ASI sensory neurons (Fig 3A-B). Strong GFP expression of the *crh-1*p*4*::*gfp* transgene was detected exclusively in the ASI neurons throughout larval development (Fig 3A-B). Expression of wild-type *crh-1* cDNA sequences driven by *crh-1*p*1* or *crh-1*p*4* promoters were both sufficient to suppress L2d formation of *crh-1* mutants (Fig 3C), suggesting that *crh-1* acts in the ASI sensory neurons. The morphology and dye-filling properties of the ASI neurons were not altered by the *crh-1* mutation (n > 50 each) (Fig EV8), suggesting that CRH-1 does not play a general role in ASI neuron development.

**Figure 3.**
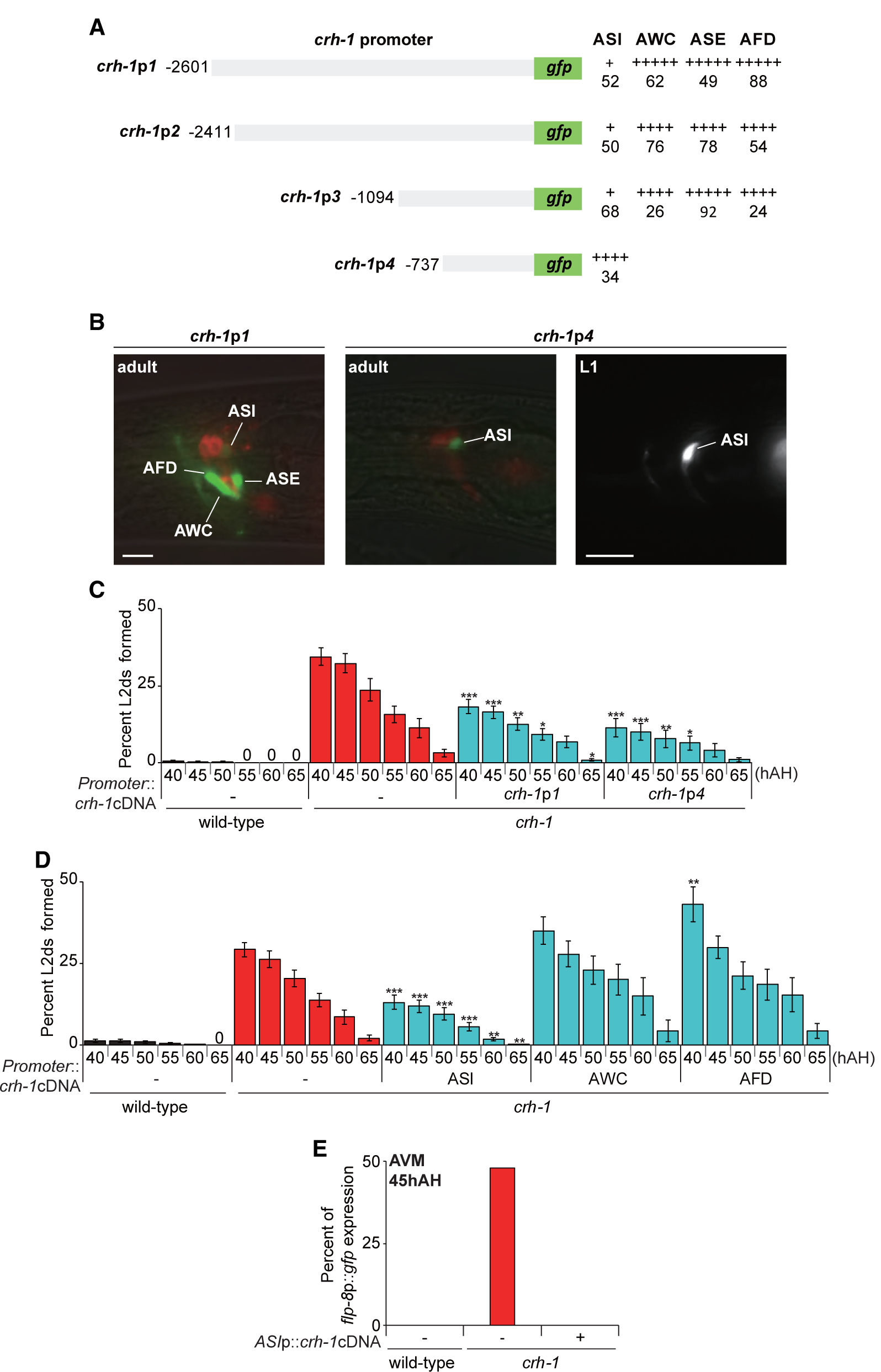
*crh-1* is expressed and acts in the ASI neurons to regulate ascr#5-mediated L2d formation. **A**. The percentage of adult transgenic animals expressing *crh-1*p*1*:: *gfp, crh-1*p*2*::*gfp, crh-1*p*3*::*gfp* or *crh-1*p*4*::*gfp* reporter construct in the indicated neurons is shown. The strength of GFP expression is indicated by the number of + symbols. At least two independent extrachromosomal lines for each construct were examined. n ≥ 60 for each. **B**. Representative images of adult or L1 larval transgenic animals expressing *gfp* expression under the control of *crh-1* promoter*1* (*crh-1*p*1*) and *crh-1* promoter*4* (*crh-1*p*4*). Adult animals were dye-filled with DiI. Scale bar: 10 µm. **C-D**. Percent of L2d formed by animals of the indicated genotypes when grown in the presence of live OP50 food and ascr#5 pheromone. N ≥ 6 for each. Error bars indicate SEM. *\*, \*\**, and *\*\*\** indicate different from *crh-1* mutants at p < 0.05, p < 0.01, and p < 0.001, respectively (one-way ANOVA with Bonferroni’s post hoc tests). **E**. Percent of *flp-8*p::*gfp* expression in the AVM neurons by animals of the indicated genotypes when grown in the presence of live OP50 food and ascr#5 pheromone. n ≥ 45 for each.

To confirm these findings, we next rescued *crh-1* mutant phenotypes by expressing *crh-1* cDNA under the control of cell-specific promoters, including *srg-47* (ASI), *ceh-36*Δ (AWC), and *ttx-1* (AFD) (Kimura et al., 2002). Whereas expression of *crh-1* in single AWC or AFD neurons did not rescue the L2d formation defects of *crh-1* mutants, ASI-specific expression rescued the phenotype to the same extent as expression driven under the endogenous *crh-1* promoter (Fig 3D). ASI-specific expression of *crh-1* also fully rescued the *flp-8* expression defects of *crh-1* mutants (Fig 3E). Taken together, we conclude that the *crh-1* gene is expressed in and functions in the ASI neurons to regulate L2d formation.

### *daf-7* TGF-β expression is decreased in the ASI neurons of *crh-1* mutants

Identification of the ASI neurons as the site of action for CRH-1 prompted us to investigate whether CRH-1 also plays a role in regulating the expression of the *daf-7* transforming growth factor-β (TGF-β), which is expressed mainly in ASI, and which plays a major role in dauer formation (Ren et al., 1996, Schackwitz et al., 1996). A feasible hypothesis for the inappropriate entry of *crh-1* mutants into the L2d stage is that the expression of *daf-7* is downregulated in *crh-1* mutants under non-dauer-inducing conditions. Previously, the *daf-7* expression has been monitored using either an integrated *daf-7*p::*gfp* reporter gene, or by single-molecule fluorescence *in situ* hybridization (smFISH) (Meisel et al., 2014, Nolan, Sarafi-Reinach et al., 2002). We found that the *daf-7*p::*gfp* reporter exhibited decreased expression in the ASI neurons of *crh-1* mutants (Fig 4A-B). The ASI-specific expression of *crh-1* restored *daf-7*p::*gfp* expression in *crh-1* mutants (Fig 4B). We next used smFISH to accurately examine the endogenous transcriptional activity of *daf-7* and observed that the fluorescence intensity of *daf-7* FISH probes in the ASI neurons was also decreased in *crh-1* (Fig 4A, C). Moreover, ascr#5 further decreased *daf-7* transcriptional activity (Fig 4C). These results demonstrate that CRH-1 is necessary to maintain a robust expression of *daf-7* in the ASI neurons.

**Figure 4.**
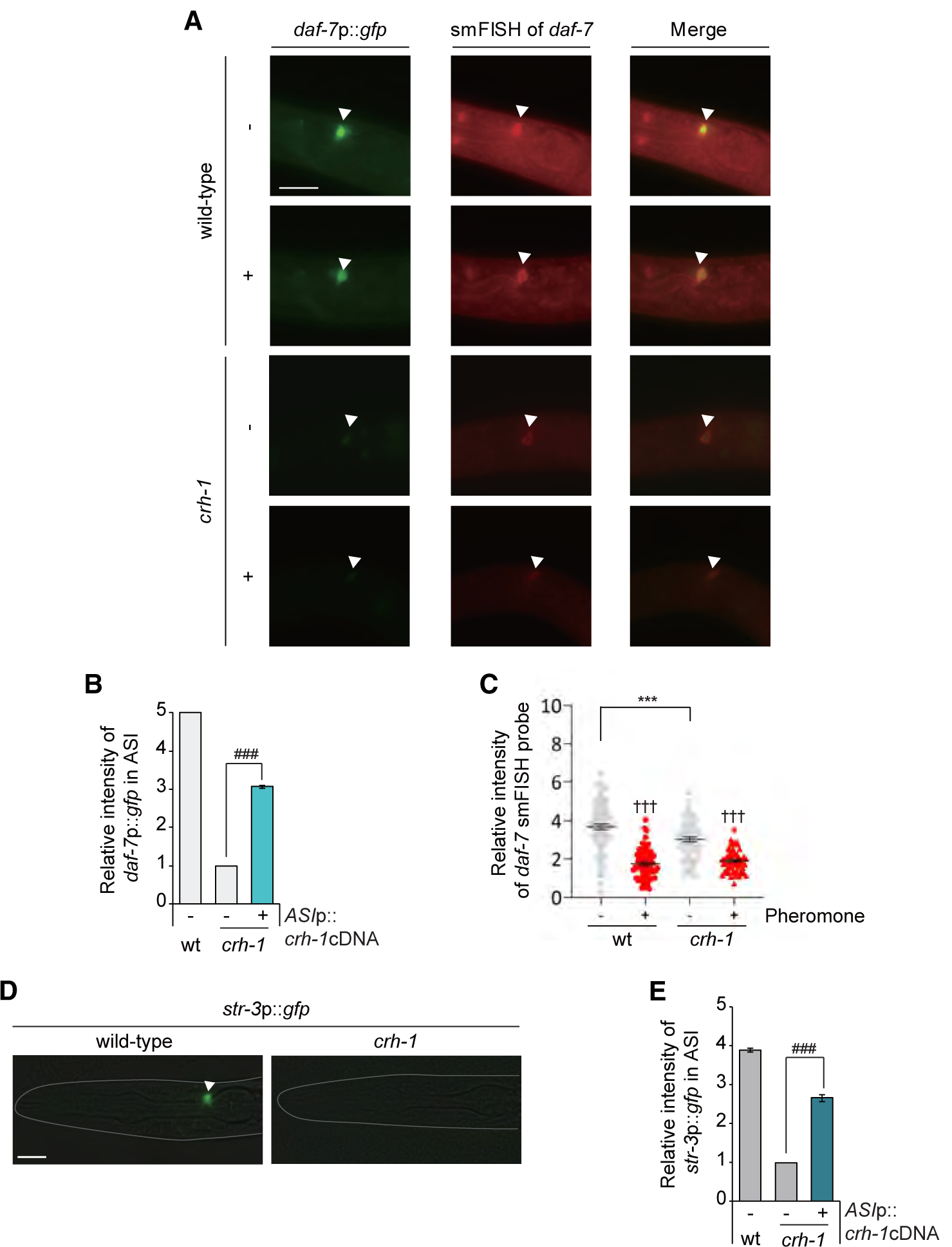
*crh-1* mutants exhibit a decrease of *daf-7* expression in the ASI neurons. **A**. Representative images of L1 larval wild-type or *crh-1* mutant animals expressing *daf-7*p::*gfp* transgene in the presence (+) or absence (-) of ascr#5. Shown is GFP expression or *daf-7* smFISH signal in the ASI neurons. Anterior is to the left. Scale bar: 50 µm. Arrowheads indicate the ASI neurons. **B**. The relative level of *daf-7*p::*gfp* fluorescence in the ASI neurons of L1 larval animals of the indicated genotypes in the absence (-) of ascr#5. **C**. Scatter plot of fluorescence intensity of *daf-7* smFISH signal in the ASI neurons of wild-type or *crh-1* mutant animals in the presence (+) or absence (-) of ascr#5. The median is indicated by a horizontal line. Each dot is the fluorescence intensity of a single neuron; n ≥ 30 neurons total each, at least two independent experiments. **D**. Representative images of L1 larval wild-type or *crh-1* mutant animals expressing *str-3*p::*gfp* in the ASI neurons. Scale bar: 10 µm. An arrowhead indicates the ASI neurons **E**. The relative level of *str-3*p::*gfp* fluorescence in the ASI neurons of L1 larval animals of the indicated genotypes in the absence (-) of ascr#5. n ≥ 30 for each. Error bars indicate SEM. ***, ^†††^, or ^###^ indicates different from wild-type, absence of ascr#5, or *crh-1* mutants at p < 0.001, respectively (student t-test).

Similar to *daf-7* expression, the expression of a candidate *str-3* chemoreceptor gene in the ASI neurons is regulated by exposure to pheromones (Kim et al., 2009, Nolan et al., 2002). Expression of an *str-3*p::*gfp* reporter gene was strongly decreased in *crh-1* mutants. This gene expression phenotype was again rescued by ASI-specific expression of *crh-1* (Fig 4D-E). Taken together, these results are consistent with the hypothesis that CRH-1 mediates transcriptional regulation of *daf-7* TGF-β and STR-3 GPCR gene expression in the ASI neurons.

### *daf-7* TGF-β signaling is required for the ascr#5-mediated L2d formation of *crh-1* mutants

*daf-7* null mutants form dauers constitutively (Daf-c) regardless of environmental conditions, although other conditions and mutants in which *daf-7* expression is perturbed result in less penetrant dauer-related phenotypes (Ren et al., 1996). We, therefore, investigated whether reduced *daf-7* expression plays a role in the inappropriate induction of L2d larvae in *crh-1* mutants. As expected, 100% of *daf-7* mutants formed L2d with abundant live OP50 food and high concentrations of ascr#5 (Fig 5A). Compared to the transient L2d formation of *crh-1* mutants, *daf-7* L2d larvae committed to dauer entry. In addition, *flp-8* expression in the AVM neurons of *daf-7* mutants initiated at 40 hAH and lasted throughout the L2d and dauer stages (Fig 5B). To confirm that DAF-7 acts downstream to CRH-1 in regulating dauer formation, we overexpressed DAF-7 specifically in the ASI neurons of *crh-1* mutants and found that it suppressed the L2d formation of *crh-1* mutants (Fig 5C). Moreover, DAF-7 expression in the ASI neurons rescued the L2d formation phenotype of *daf-7* mutants (Fig 5D), further supporting a role of the ASI neurons in the L2d formation of *crh-1* mutants.

**Figure 5.**
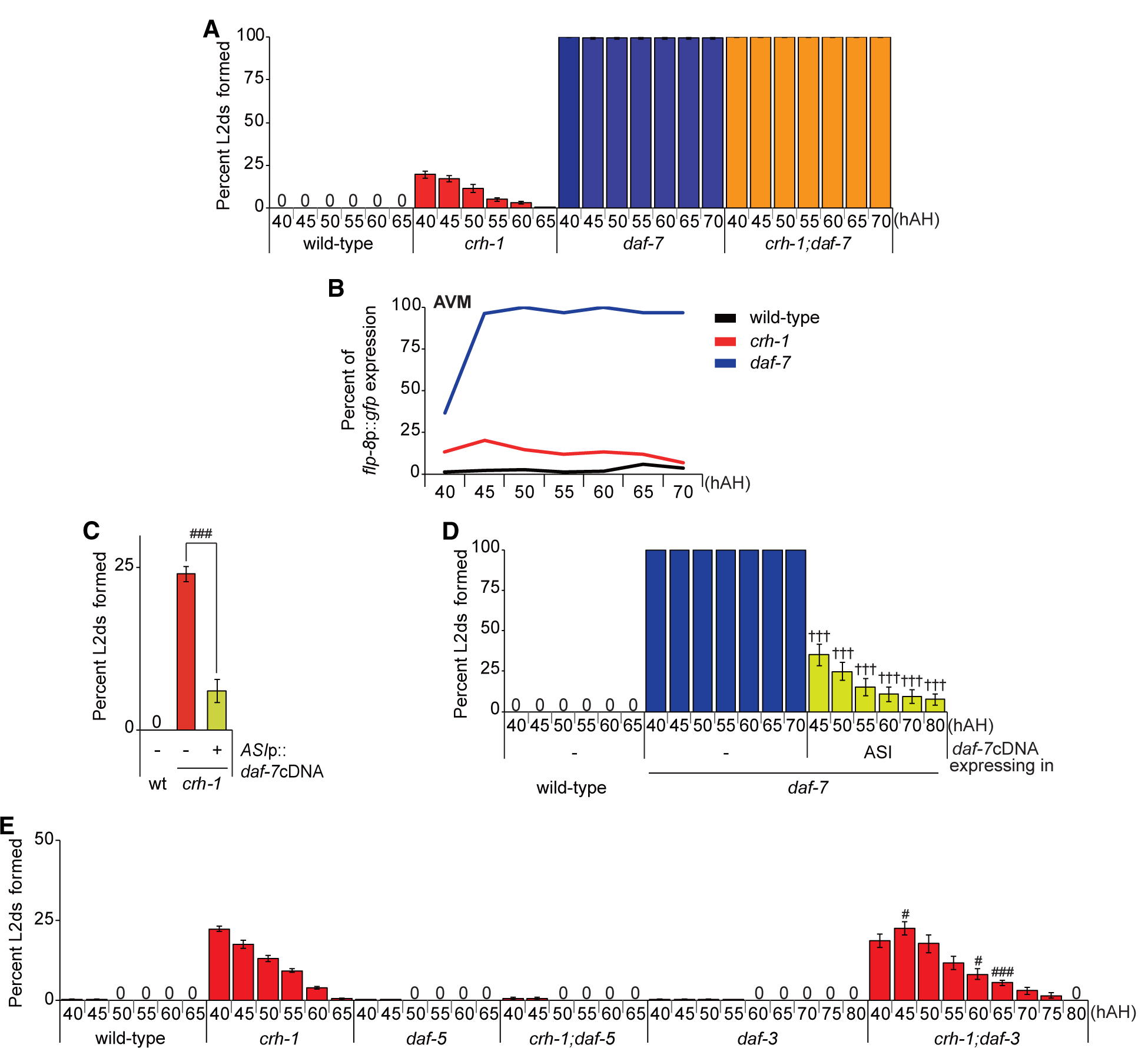
*daf-7* mediates the ascr#5-mediated L2d formation of *crh-1* mutants. **A**. Percent of L2d formed by animals of the indicated genotypes when grown in the presence of live OP50 food and ascr#5 pheromone. N ≥ 4 for each. **B**. Percent of *flp-8*p::*gfp* expression in the AVM neurons by animals of the indicated genotypes when grown in the presence of live OP50 food and ascr#5 pheromone. n ≥ 30 for each. **C-E**. Percent of L2d formed by animals of the indicated genotypes when grown in the presence of live OP50 food and ascr#5 pheromone. N ≥ 8 (**C**), 6 (**D**), 4 (**E**) for each. Error bars represent the SEM. ^###^ or ^†††^ indicates different from *crh-1* or *daf-7* mutants at p < 0.001, respectively (student t-test).

It was previously shown that the Daf-c phenotype of *daf-7* mutants was suppressed by loss-of-function mutations of the *daf-3* SMAD or *daf-5* SNO/SKI transcription factor genes (da Graca, Zimmerman et al., 2004, Patterson, Koweek et al., 1997), whereas mutation of *daf-16* represses Daf-c mutations in the insulin signaling pathway (Kimura et al., 1997). We found that the loss of *daf-5* function fully suppressed the L2d phenotype of *crh-1* mutants, whereas the loss of *daf-3* or *daf-16* function did not affect the L2d formation of *crh-1* mutants (Fig 5C, Fig EV9). These results suggest that *daf-5* acts downstream of *crh-1* in ascr#5-mediated L2d formation and that the *daf-7* TGF-β signaling pathway mediates the L2d formation of *crh-1* mutants.

### CRH-1 directly regulates *daf-7* expression via a conserved CRE motif

Like mammalian CREB, CRH-1 can act as a sequence-specific transcription factor to regulate gene expression via cAMP-response elements (CRE; TGACGTCA) (Comb, Birnberg et al., 1986) (Fig 6A). To determine whether CRH-1 directly regulates the expression of *daf-7* in the ASI neurons, we first analyzed the promoter regions of the *daf-7* gene. Although we could not identify any DNA sequences which showed exact matches to the CRE site within 3.1 kb upstream of the *daf-7* initiator codon, we identified ten DNA sequences that contain the consensus ACGT CRE core sequence within this sequence (Fig 6B). We mutated these sequences in the context of the *daf-7*p::*gfp* reporter gene, and found that mutations in one specific sequence, “GTACGTAC” located ∼2380 bp upstream of the translational start site caused decreased *daf-7*p::*gfp* expression in the ASI neurons (Fig 6B-C). This mutation did not affect the faint reporter expression we observed in ASJ neurons (Fig EV10), indicating the specificity of this motif in the regulation of ASI expression of *daf-7* (Meisel et al., 2014). Moreover, the decreased ASI *daf-7* expression by mutations of the CRE site was not altered in *crh-1* mutant adults (Fig 6D). These results support our hypothesis that CRH-1 binds this putative CRE in the *daf-7* promoter to regulate its expression in ASI neurons.

**Figure 6.**
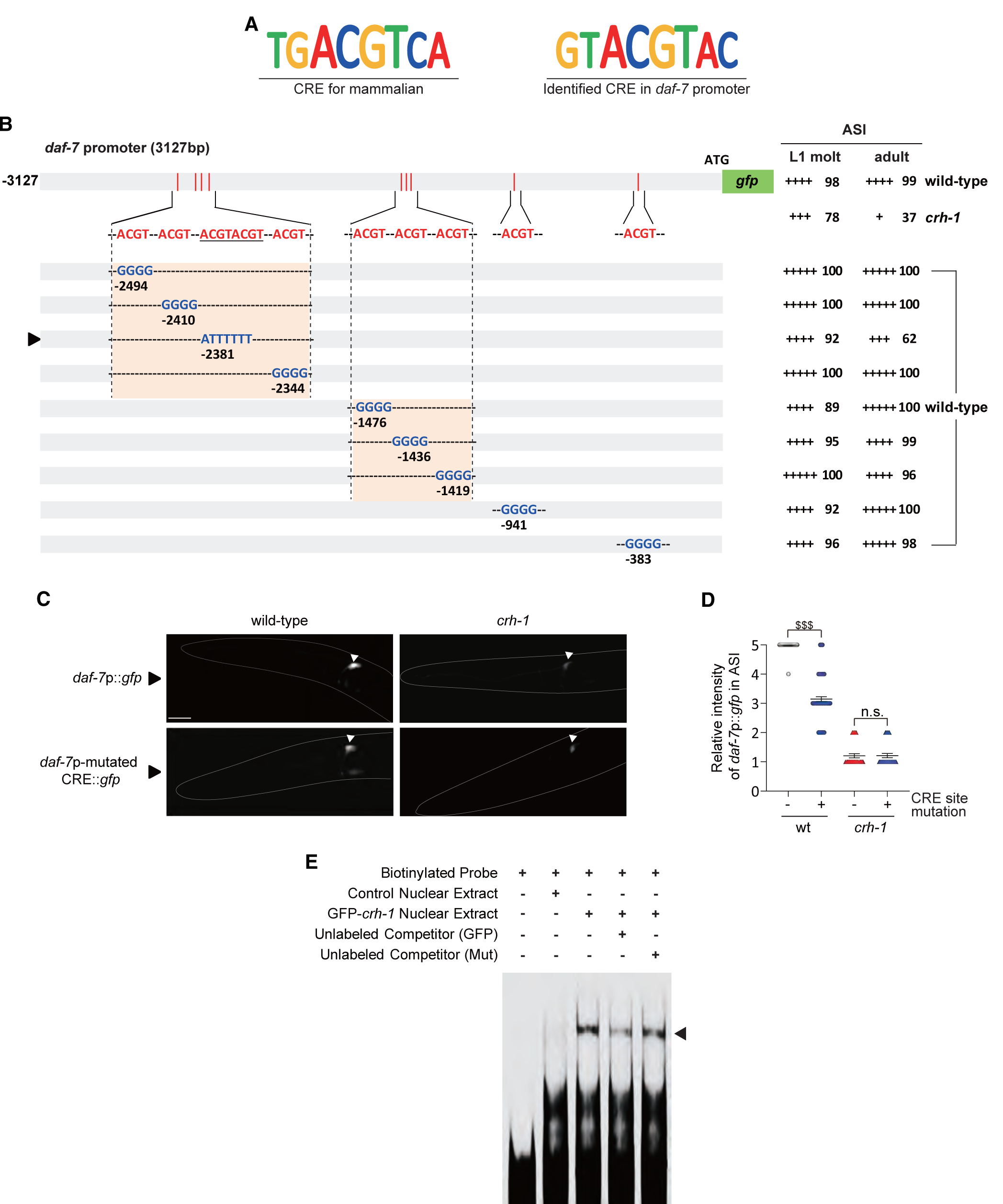
CRH-1 directly binds a CRE motif in the *daf-7* promoter. **A**. Shown are the conserved mammalian CRE motif and the newly identified CRE motif in the *daf-7* promoter. **B**. The percentage of transgenic animals expressing *daf-7*p:: *gfp* reporter construct in the ASI neurons is shown. GFP fluorescence was observed either in wild-type or *crh-1* mutant animals. The Strength of GFP expression is indicated by the number of + symbols. Wild-type nucleotides are indicated in red, mutated nucleotides in blue. An arrowhead indicates *daf-7*p-mutated CRE promoter. At least two independent extrachromosomal lines for each construct were examined. N ≥ 30 for each. **C**. Representative images of L1 larval wild-type or *crh-1* mutant animals expressing *daf-7*p::*gfp* or *daf-7*p-mutated CRE::*gfp* in the ASI neurons. Scale bar: 50 µm. **D**. Scatter plot of fluorescence intensity of *daf-7*p::*gfp* fluorescence in the ASI neurons of adult wild-type or *crh-1* mutant animals with (+) or without (-) of CRE site mutation. The median is indicated by a horizontal line. Each dot is the fluorescence intensity of a single neuron; n ≥ 30 neurons total each, at least two independent experiments. Error bars represent the SEM. ^$$$^ indicates different at p < 0.001, respectively (student t-test). **E**. Nuclear extracts isolated from control (2^nd^ lane) or GFP-*crh-1* (3^rd^, 4^th^, 5^th^ lane) transfected cells were analyzed by EMSA using biotinylated *daf-7*p-CRE probes (2^nd^, 3^rd^, 4^th^, 5^th^ lane) with non-biotinylated *daf-7*p-WT-CRE (4^th^ lane) and *daf-7*p-Mut-CRE (5^th^ lane) competitors as indicated. An arrowhead indicates a complex of CRE probes and CRH-1.

To confirm the direct binding of CRH-1 to this CRE motif *in vitro*, we performed electrophoretic mobility shift (EMSA) assays. Specific CRE binding activity was evident in GFP-CRH-1 overexpressing nuclei, and absent in control nuclear extract (Fig 6E). Furthermore, this CRE binding activity was significantly diminished by co-incubation with wild-type non-biotinylated probes, whereas mutant unlabeled probes failed to effectively compete (Fig 6E). These results indicate that CRH-1 can directly bind to the CRE motif identified in the *daf-7* promoter sequence, which, in turn, may enhance *daf-7* expression in the ASI neurons.

Thus, we have identified CRH-1 as a critical mediator of ASI state in developing *C. elegans*. CRH-1 promotes the expression of *daf-7* in the presence of pheromone and abundant food to suppress inappropriate passage through L2d, which ultimately delays reproductive maturation of the animal and may be disadvantageous to fitness.

## Discussion

Dauer entry in *C. elegans* is gated by sequential developmental decisions, first, at L1 to transition to either L2 or L2d, and second, a commitment to either diapause or reproductive growth (Fig 1A). Precociously or inadvertently entering dauer, has the potential to carry a large fitness cost for an animal, given the rapid development and fecundity of *C. elegans*. Here we have shown that the CREB homolog CRH-1 is a key regulator of the first decision, whether or not to enter L2d. *crh-1* mutants inappropriately express the L2d marker gene *flp-8* in ALM, AVM, and PLM neurons and transiently form L2d larvae in non-dauer-inducing conditions. However, *crh-1* mutants do not commit to dauer entry in the absence of additional environmental cues and thus ultimately make the correct developmental decision. Our data support a model in which CRH-1 positively regulates dauer-inhibiting *daf-7* TGF-β expression in the ascr#5-sensing ASI neurons (Fig 7). The loss of *crh-1* function leads to decreased *daf-7* TGF-β expression, which provides a sensitized genetic background for further ascr#5-mediated *daf-7* TGF-β repression via SRG-36/37 GPCRs (Fig 7). When the *daf-7* expression is eliminated, animals constitutively enter the dauer state, and thus *daf-7* expression serves as a critical rheostat for the determination of *C. elegans* developmental trajectory (Kim et al., 2009, Ren et al., 1996, Schackwitz et al., 1996). Our results demonstrate that worms exploit distinct strategy and molecules at different dauer decision points and CRH-1 acts in ASI and gates the initial L1-to-L2d decision in response to pro-growth environmental signals (food), prior to dauer commitment, and thus *crh-1* mutant provides a unique opportunity to study the under-explored L1 to L2/L2d transition.

**Figure 7.**
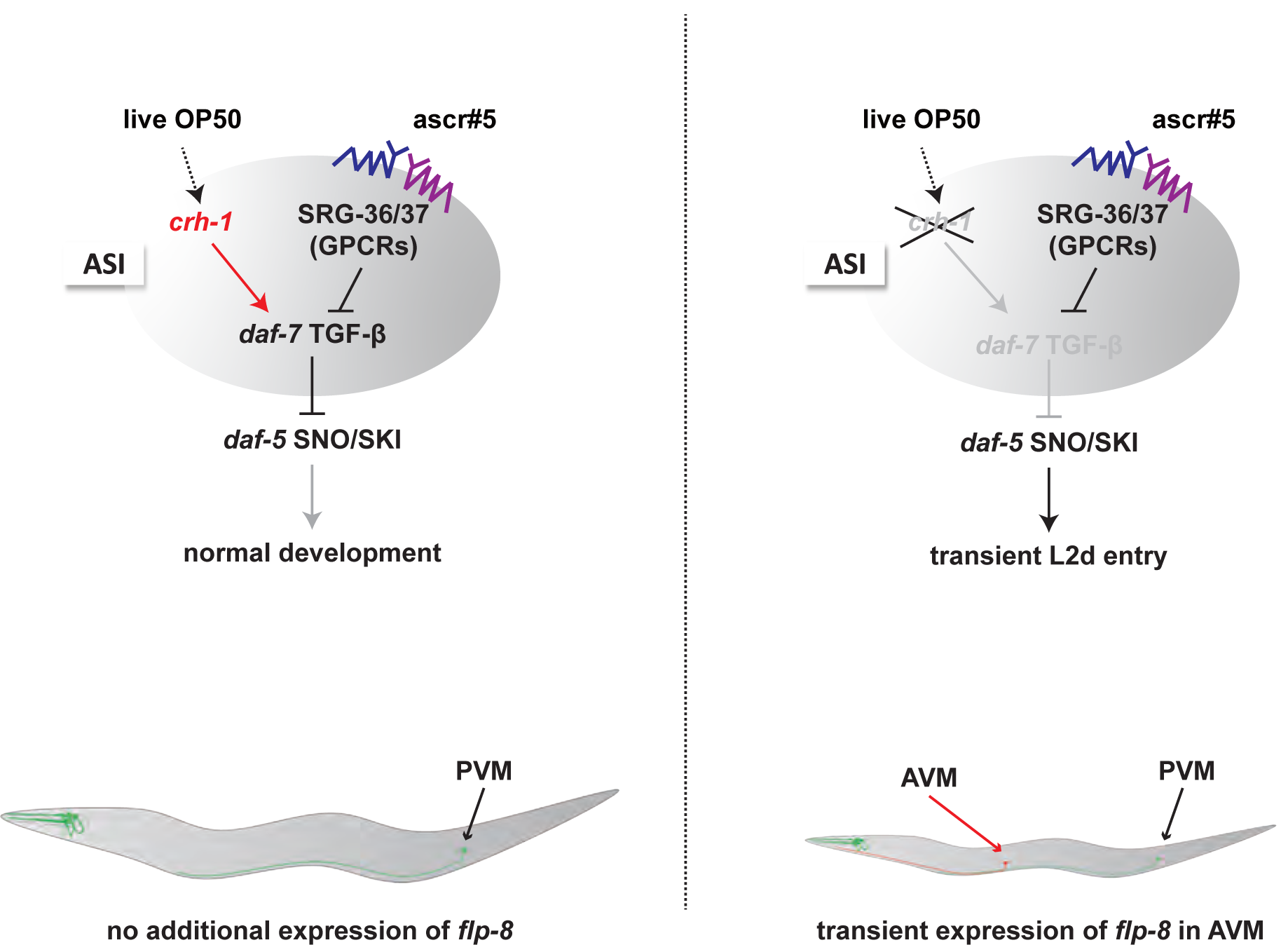
A model of the role of CRH-1 in ascr#5-mediated L2d formation. CRH-1 positively regulates dauer-inhibiting *daf-7* TGF-β expression in the ascr#5-sensing ASI neurons. In *crh-1* mutants, the decreased *daf-7* TGF-β expression provides a sensitized genetic background for further ascr#5-mediated *daf-7* TGF-β repression via *srg-36/37* GPCRs which leads to transient L2d formation and *flp-8* expression of AAP-TRN.

We have shown that an abundance of live OP50 food and high concentrations of ascr#5 do not induce dauer formation (L2d to dauer transition) but do induce L2d formation (L1 to L2d transition) in *crh-1* mutants. These results suggest CRH-1 is necessary to give the appropriate weight to pro-growth food cues in the environment such that the presence of pheromone in the environment is sufficient to promote L2d formation but not dauer commitment (Golden & Riddle, 1984). Moreover, ascr#5 but not ascr#2 or ascr#3 causes L2d formation in *crh-1* mutants, indicating that specific, but not all, pheromone component(s) may affect the decision for L1 to transition to L2 or L2d and worms weigh pheromone signals differently at different dauer decision points. ascr#5 is the most potent component of the dauer inducing ascaroside pheromones (Butcher et al., 2008) and is produced in higher amounts in well-fed worms than in starved worms, indicating that ascr#5 may be a *bona fide* indicator of healthy populations or favorable environments (von Reuss et al., 2012). Despite this hypothesis, very high concentrations of ascr#5 can elicit L2d formation in wild-type animals, indicating that *crh-1* mutants are precocious in L2d entry. Given that ascr#5 further reduces *daf-7* expression in the ASI neurons of *crh-1* mutants, we infer that CRH-1 acts as a food sensor to set the basal level of *daf-7* expression. Furthermore, the two-step process to enter dauer likely allows animals to be more flexible and adjust to rapidly changing environments (Avery, 2014).

Previously, a large number of genes have been shown to be differentially expressed in dauer larvae (Lee, Shih et al., 2017, Wang & Kim, 2003). Interestingly, expression of most, but not all, *flp* genes, including *flp-8* are up-regulated during dauer development (Lee et al., 2017), and several *flp* genes have shown to be involved in dauer development and behaviors (Cohen, Reale et al., 2009, Lee et al., 2017). Since the *flp-8* expression in the AAP-TRN begins at the L2d stage and is maintained throughout the dauer stage, the function of FLP-8 neuropeptide in the AAP-TRN of L2d and dauer larvae is intriguing. Although the exact function of FLP-8 has not been determined, a family of FLP neuropeptides appears to inhibit overall circuit activity (Li & Kim, 2014). Previously, Chen and Chalfie (Chen & Chalfie, 2014) showed that dauer larvae exhibited reduced touch sensitivity. Thus, it is possible that FLP-8 expression in the AAP-TRN could inhibit circuit activity underlying touch sensation, which leads to decreased mechanosensation in dauer and likely L2d larvae. It would be the next question to be explored in the future.

Transforming growth factor β (TGF-β) signaling mediates diverse physiological roles, including cell differentiation, morphogenesis, tissue homeostasis, regeneration, and immune response, while the misregulation of TGF-β signaling promotes distinct disease states such as cancer progression (see review by Massague (Massague, 2008)). While cellular response pathways to secretion and diffusion/trafficking of the TGF-β ligand are well-characterized, relatively little is known about the regulation of its expression (Kim, Jeang et al., 1989). In *C. elegans*, TGF-β signaling regulates body size, innate immunity, aging, cell fate specification, differentiation, and dauer formation (see review by Savage and Padgett (Savage-Dunn & Padgett, 2017)). Although the downstream components of TGF-β signaling are evolutionarily conserved and well-characterized, upstream transcription factors and *cis*-regulatory elements by which expression of TGF-β genes is controlled are not as well characterized. Here, we have shown that the CREB homolog CRH-1 directly binds a novel CRE in the *daf-7* TGF-β promoter, which must integrate multiple external and internal signals and positively regulates its expression in the ASI neuroendocrine cells. Future studies are needed to identify potential CRH-1 co-factors, which may help to refine or modulate its actions at the *daf-7* promoter. Similar to mammalian CREB, *C. elegans* CRH-1 functions in many developmental and behavioral processes, including longevity, learning and memory, and now, developmental plasticity. Thus, CREB acts as a determinant of a context-dependent role of TGF-β signaling and provides a useful model to study TGF-β gene expression.

## Materials and Methods

### Strains and constructs

The N2 Bristol strain was used as wild-type. All strains were maintained and grown on NGM agar plates seeded with *E. coli* OP50 at 20°C (Brenner, 1974). All used strains in this study are listed in Table EV1. Promoters and cDNA used for rescue experiments of *crh-1* and *daf-7* were described in Park et al. (Park, Choi et al., 2019). The *daf-7* promoter (3127bp) for expression and CRE site analysis was produced by polymerase chain reactions using forward and reverse primers (see Table EV2) and subcloned into a pPD95.77 vector. All plasmids were injected at 0.1 ng (*flp-8* expression analysis), 10 ng (rescue experiments), or 50 ng (expression analysis) with *unc-122*p::*dsRed* as an injection marker.

### Pheromone preparation and assay plates

Crude pheromone was prepared following the protocol described by Golden and Riddle (1984). Before assay, each crude pheromone batch was tested for its efficiency to induce dauer formation. The assay plates containing crude pheromone were prepared by spreading pheromone and drying for 3-4 h. The ascaroside pheromone components were chemically synthesized following Butcher et al. (Butcher et al., 2007, Butcher et al., 2008). Before use, pheromone was diluted with dH_2_O from a 3 mM stock solution of pheromone in 100% ethanol. For the assay plates containing synthetic ascarosides, the pheromone solution was mixed with agar solution before pouring.

### L2d formation Assay

Synchronized adult animals were placed onto 3.5 cm assay plates seeded with live OP50 and containing ethanol or pheromone at 25°C and allowed to lay eggs for 1-2 h. Then, adult animals were removed, and assay plates were placed at 25°C for 40 h before observing the morphology of worms or marker gene expression and counting as L2d larvae.

### Dauer formation Assay

A dauer formation assay was performed as previously described by Golden and Riddle (Golden & Riddle, 1984) and Neal et al. (Neal, Kim et al., 2013). Synchronized adult animals were allowed to lay eggs at 25°C for 4-5 h on 3.5 cm assay plates. After the plates contained approximately 65-85 eggs/plate, adult animals were removed, and the plates were placed at 25°C for 68-72 h. Dauer and non-dauer animals were identified by visual inspection. Assay plates were made with Noble agar (BD Biosciences) lacking peptone. Heat-killed OP50 bacteria were seeded on each plate. For each experiment, all strains were assayed in parallel in at least four independent experiments.

### *daf-7* smFISH

*daf-7* smFISH was performed as previously described by Meisel et al. (Meisel et al., 2014) by using the Stellaris smFISH fluorescent probe and buffer sets (Biosearch). Worms were washed in 1XPBS and fixed in 3.7% formaldehyde in PBS for 45 min at 4°C and rinsed twice with 1XPBS. Then, worms were immersed in 70% ethanol for 24 h at 4°C and washed by buffer A (Biosearch) for 5 minutes at room temperature. Then, a hybridization buffer containing the *daf-7* probe (125 nM, Biosearch) was added and incubated overnight at 30°C. Worms were washed twice with buffer A. Then, worms were mounted onto glass slides by mounting medium (Sigma). All smFISH images were acquired by a Zeiss Axio Imager using 63x objectives and a CCD camera (Hamamatsu). Quantification was performed using ImageJ software to extract mean intensity and regions of interest. The relative mean intensity of fluorescence was normalized by area.

### Quantification of GFP expression

For GFP quantification, the worms were anesthetized in sodium azide on an agar pad, and GFP fluorescence was observed with a Zeiss Axio Imager using 40x (for the adult stage) and 63x (for L1) objectives and a CCD camera (Hamamatsu). The relative expression level of GFP was measured at each developmental stage. The relative GFP levels were rated from 1 (dim) to 5 (bright) by visual inspection, and these values were confirmed using Image J software.

### SEM (Scanning Electron Microscope) imaging

Samples were prepared by critical point dryer (Leica EM CPD300) after sequential ethanol dehydration. Dried samples were coated with gold using a sputter coater and then were imaged with a HR FE-SEM (Hitachi, SU8020, CCRF DGIST).

### Electrophoretic mobility shift assay (EMSA)

Nuclear extract preparation from HEK293T cells and in vitro EMSA reactions were carried out as previously described Kim et al. (Kim, Kempf et al., 2003), and biotinylated EMSA probes and unlabeled competitor probes were synthesized by using a LightShift® Chemiluminescent EMSA Kit (Thermo Scientific). The *daf-7*p-WT-CRE and *daf-7*p-mut-CRE probes with the following sequences were employed:

*daf-7*p-WT-CRE: 5’ attagggtacgtacgtcaatattagggtacgtacgtcaatattagggtacgtacgtcaat 3’ (60 mer)

*daf-7*p-mut-CRE: 5’ attagggtatttttttcaatattagggtatttttttcaatattagggtatttttttcaat 3’ (60 mer)

## Acknowledgements

We would like to thank the *Caenorhabditis* Genetics Center (NIH Office of Research Infrastructure Programs, P40 OD010440) and the National BioResource Project (Japan) for strains. We also would like to thank Dennis H. Kim, Seung-Jae V. Lee, Sun-Kyung Lee, and K. Kim lab members for their helpful comments and discussion on the manuscript. This work was supported by the National Research Foundation of Korea (NRF-2020R1A4A1019436, NRF-2018R1A2A3074987) (K.K.) and (NRF-2017R1E1A1A01074980) (D.K.), and NIH R01GM0 87533 (R.A.B).

## Author contributions

JP, HO, SJN, PS, DK, and KK conceived and designed the experiments. JP, HO, DYK, YC, and YJP performed the experiments. JP, HO, SJN, PS, DK, and KK analyzed the data. ADR and RAB contributed reagents. JP and KK wrote the manuscript with input from all authors.

## Expanded View Figure legends

**Figure EV1.**
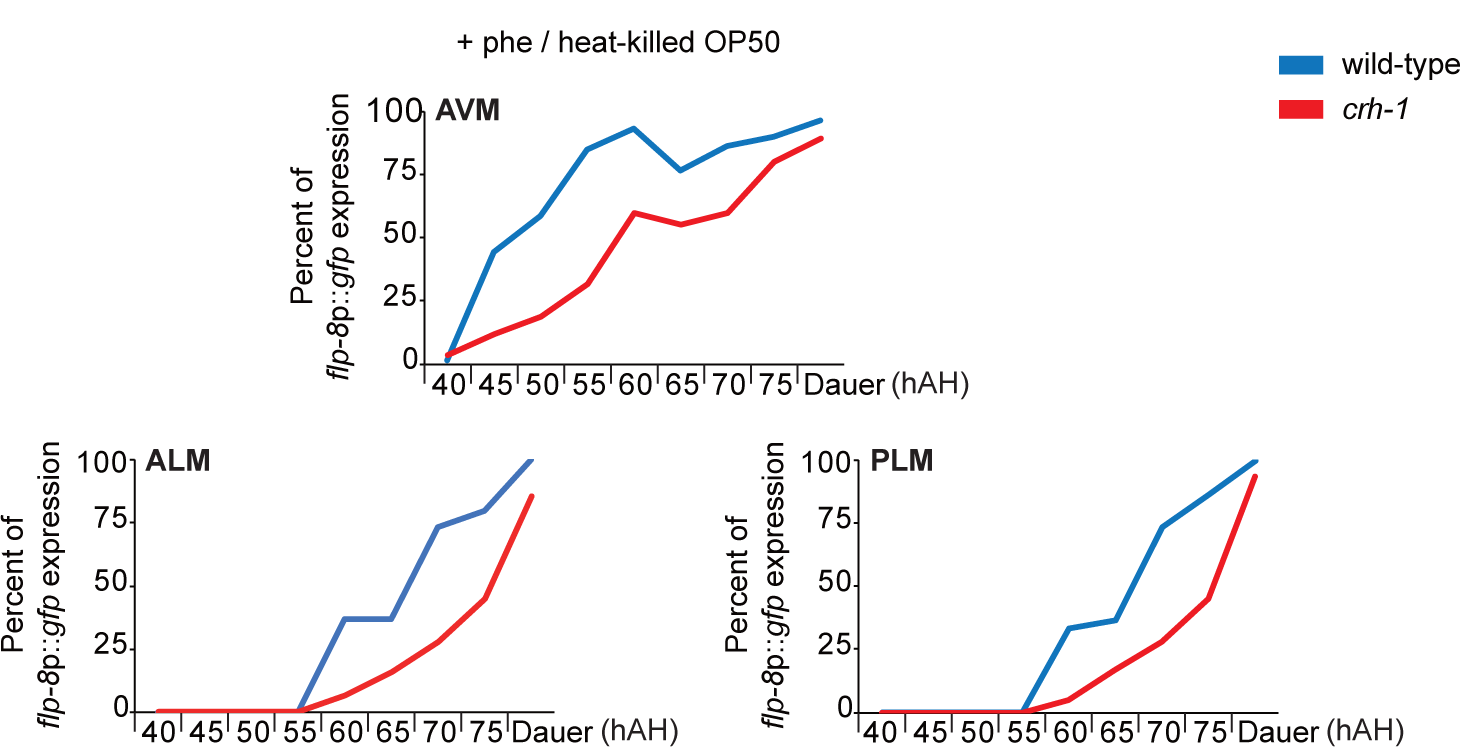
*crh-1* mutants exhibit expression of *flp-8* in the AAP-TRN in dauer-inducing conditions. Percent of *flp-8*p::*gfp* expression in the AVM, ALM and PLM neurons by wild-type or *crh-1* mutant animals when grown in the presence of heat-killed OP50 food and ascr#5 pheromone. n ≥ 30 for each. hAH, hour after hatching.

**Figure EV2.**
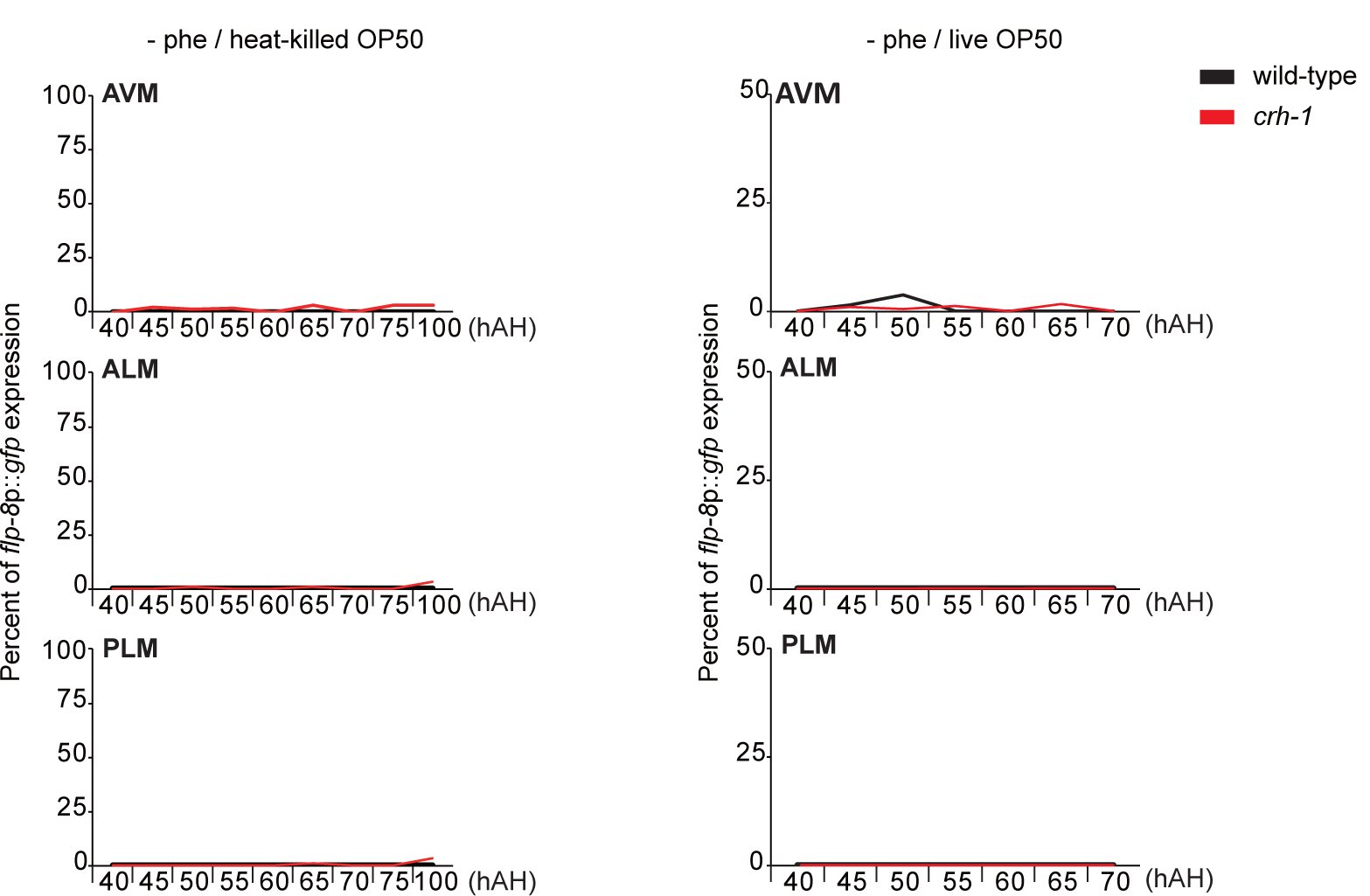
ascr#5 is required for *flp-8*p::*gfp* expression in the AAP-TRN in wild-type or *crh-1* mutant animals. Percent of *flp-8*p::*gfp* expression in the AVM, ALM and PLM neurons by wild-type or *crh-1* mutant animals when grown in the presence of heat-killed OP50 food (left) or live OP50 (right). n ≥ 30 for each. hAH, hour after hatching.

**Figure EV3.**
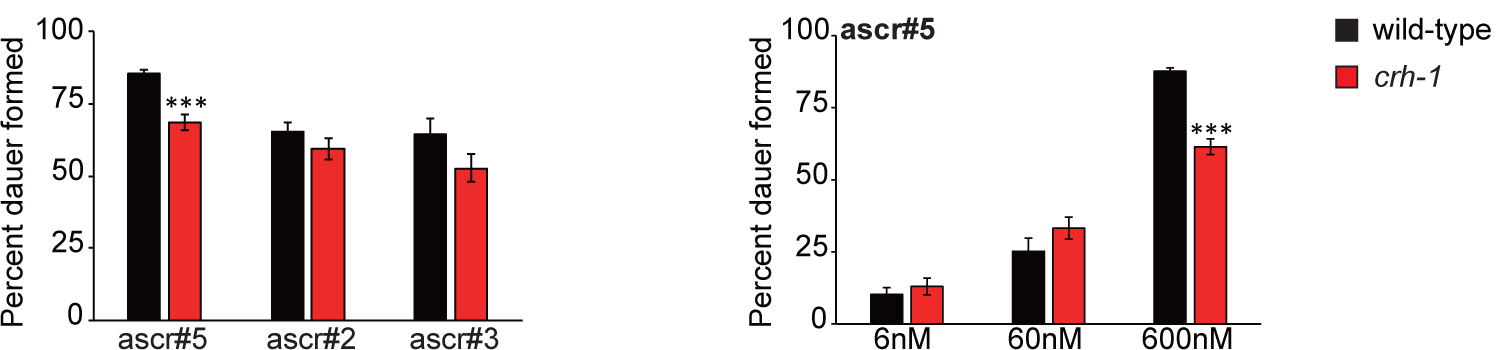
*crh-1* mutants exhibit normal or weakly decreased pheromone-mediated dauer formation. Percent of dauer formed by wild-type or *crh-1* mutant animals when grown in the presence of heat-killed OP50 food and ascr#5, ascr#2, or ascr#3 pheromone.

**Figure EV4.**
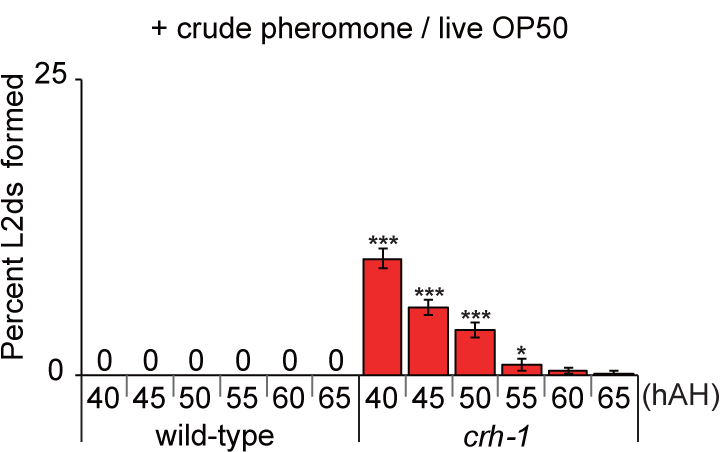
*crh-1* mutants form L2d larvae in a non-dauer inducing condition. Percent of L2d formed by wild-type or *crh-1* mutant animals when grown in the presence of live OP50 food and crude pheromone. N ≥ 5 for each. Error bars indicate SEM. *\*, \*\**, and *\*\*\** indicate different from wild-type at p < 0.05, p < 0.01, and p < 0.001, respectively (student t-test)

**Figure EV5.**
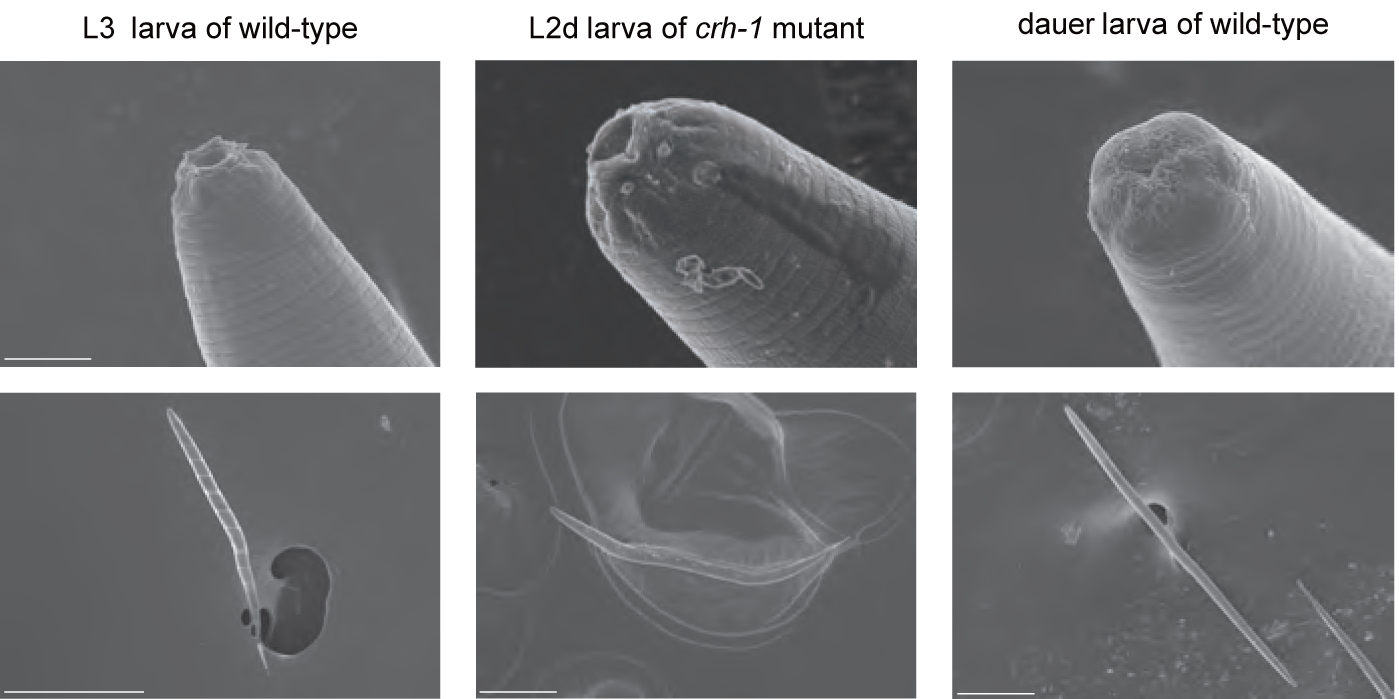
Morphology of mouths of wild-type L3 larva, *crh-1* mutant L2d larva, and wild-type dauer larva. Shown are images of scanning electron microscopy. Scale bar: 5µm (top) and 100µm (bottom).

**Figure EV6.**
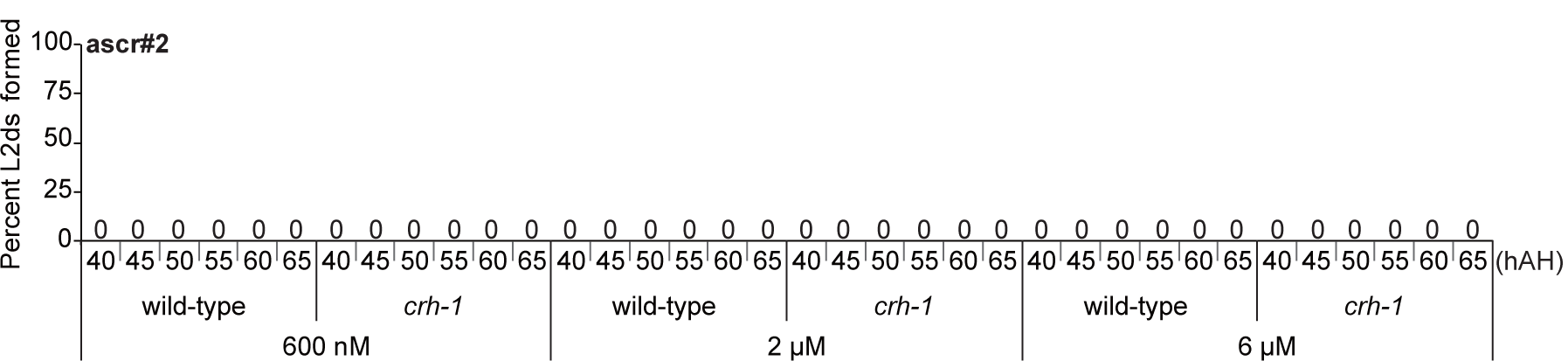
*crh-1* mutants do not form L2d larvae in the presence of ascr#2. Percent of L2d formed by wild-type or *crh-1* mutant animals when grown in the presence of live OP50 food and three different concentrations of ascr#2 pheromone. N ≥ 5 for each.

**Figure EV7.**
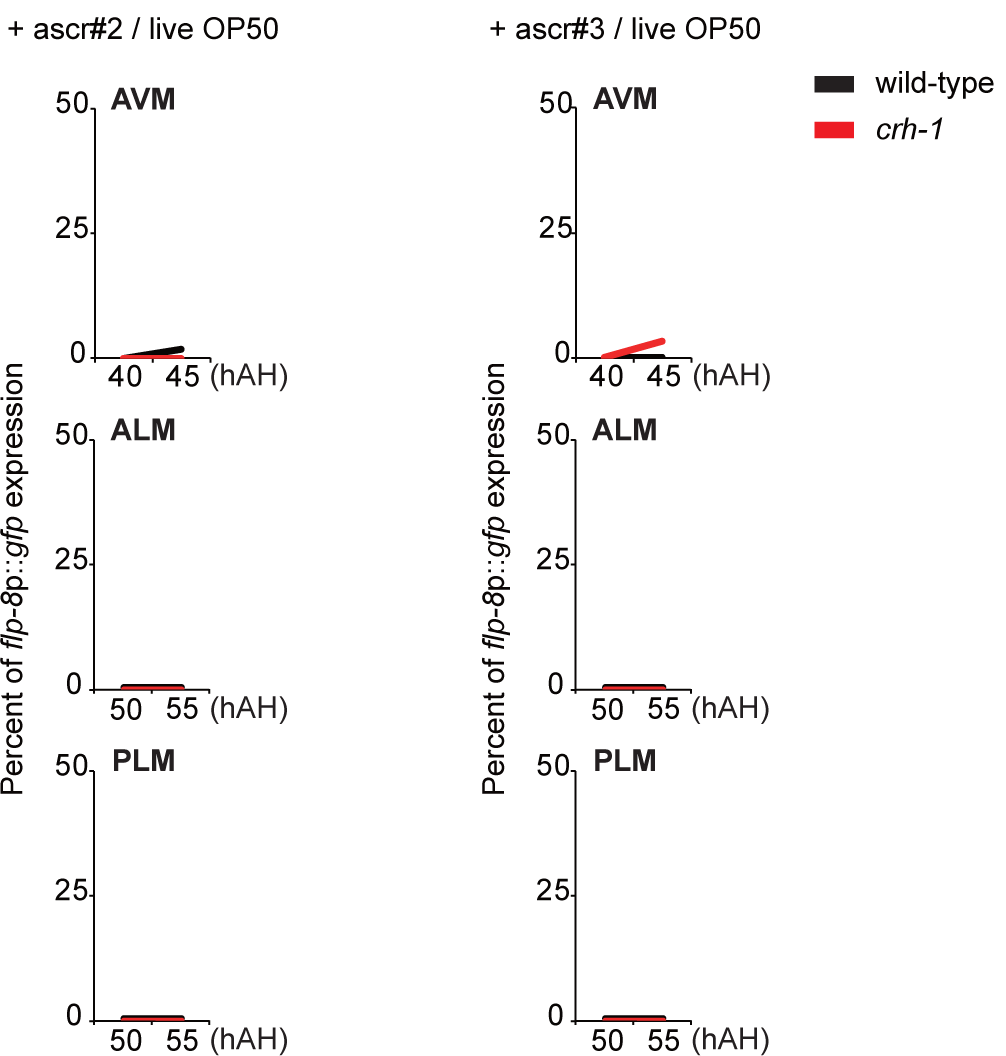
*crh-1* mutants do not exhibit *flp-8* expression in the AAP-TRN in response to ascr#2 and ascr#3. Percent of *flp-8*p::*gfp* expression in the AVM, ALM and PLM neurons by wild-type or *crh-1* mutant animals when grown in the presence of live OP50 and ascr#2 or ascr#3. n ≥ 30 for each. hAH, hour after hatching.

**Figure EV8.**
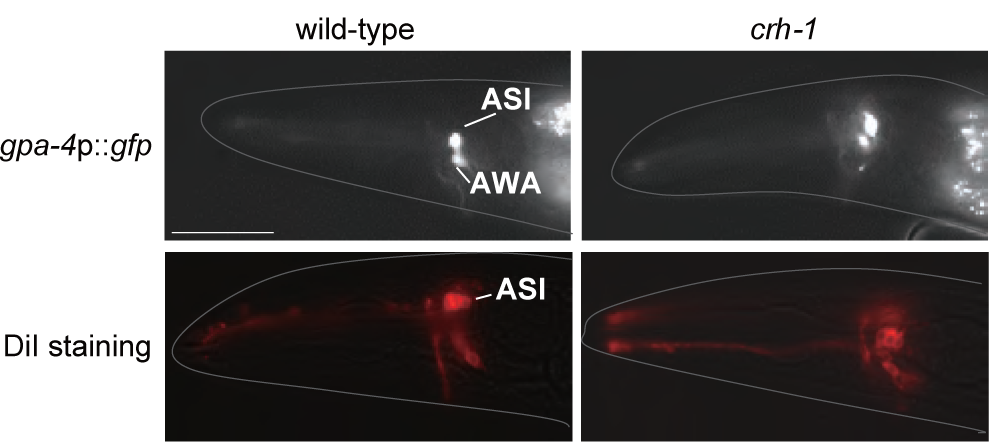
The ASI morphology and dye-filling properties are not altered in the *crh-1* mutants. Shown are images of wild-type or *crh-1* mutant animals expressing *gpa-4*p::*gfp* transgene or stained with DiI. Scale bar: 10 µm.

**Figure EV9.**
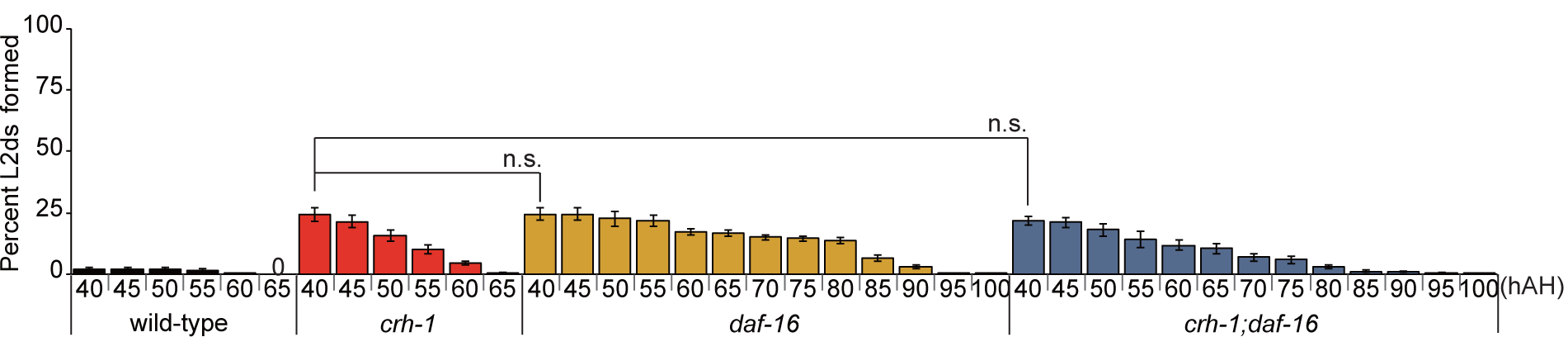
*daf-16* mutation does not affect the L2d formation of *crh-1* mutants. Percent of L2d formed by animals of the indicated genotypes when grown in the presence of live OP50 food and ascr#5 pheromone. N ≥ 4 for each. Error bars indicate SEM. N.S. not significantly different (one-way ANOVA with Bonferroni’s post hoc tests).

**Figure EV10.**
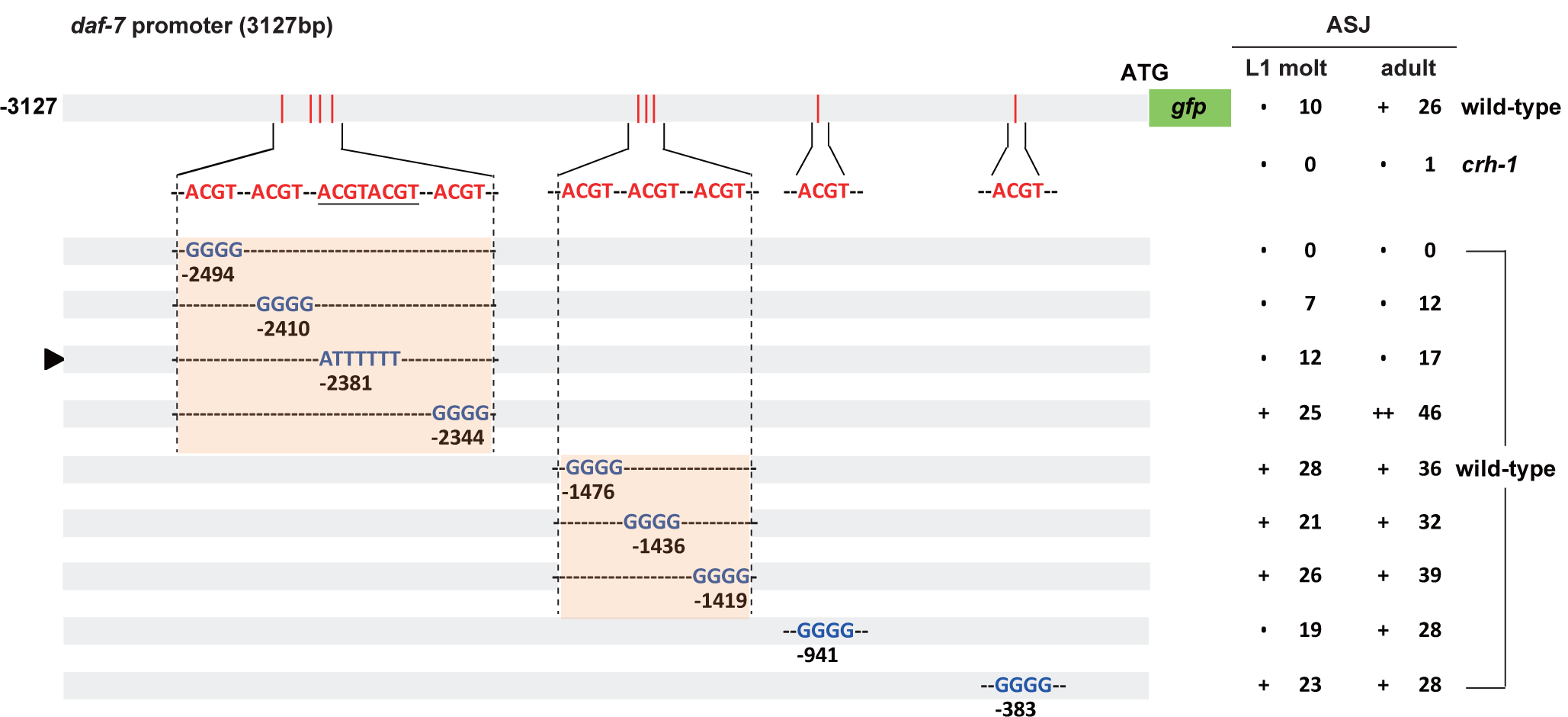
Mutation in CRE motif of *daf-7* promoter does not affect *daf-7* expression in the ASJ neurons. The percentage of transgenic animals expressing *daf-7*p::*gfp* reporter construct in the ASJ neurons is shown. GFP fluorescence was observed either in wild-type or *crh-1* mutant animals. The Strength of GFP expression is indicated by the number of + symbols. Wild-type nucleotides are indicated in red, mutated nucleotides in blue. An arrowhead indicates *daf-7*p-mutated CRE promoter. At least two independent extrachromosomal lines for each construct were examined. n ≥ 30 for each.

## References

Altarejos JY, Montminy M (2011) CREB and the CRTC co-activators: sensors for hormonal and metabolic signals. Nat Rev Mol Cell Biol 12: 141–51

Avery L (2014) A model of the effect of uncertainty on the C elegans L2/L2d decision. PLoS One 9: e100580

Brenner S (1974) The genetics of Caenorhabditis elegans. Genetics 77: 71–94

Butcher RA (2017) Small-molecule pheromones and hormones controlling nematode development. Nat Chem Biol 13: 577–586

Butcher RA, Fujita M, Schroeder FC, Clardy J (2007) Small-molecule pheromones that control dauer development in Caenorhabditis elegans. Nat Chem Biol 3: 420–2

Butcher RA, Ragains JR, Kim E, Clardy J (2008) A potent dauer pheromone component in Caenorhabditis elegans that acts synergistically with other components. Proc Natl Acad Sci U S A 105: 14288–92

Cassada RC, Russell RL (1975) The dauerlarva, a post-embryonic developmental variant of the nematode Caenorhabditis elegans. Dev Biol 46: 326–42

Chen X, Chalfie M (2014) Modulation of C. elegans touch sensitivity is integrated at multiple levels. J Neurosci 34: 6522–36

Chen YC, Chen HJ, Tseng WC, Hsu JM, Huang TT, Chen CH, Pan CL (2016) A C. elegans Thermosensory Circuit Regulates Longevity through crh-1/CREB-Dependent flp-6 Neuropeptide Signaling. Dev Cell 39: 209–223

Cohen M, Reale V, Olofsson B, Knights A, Evans P, de Bono M (2009) Coordinated regulation of foraging and metabolism in C. elegans by RFamide neuropeptide signaling. Cell Metab 9: 375–85

Comb M, Birnberg NC, Seasholtz A, Herbert E, Goodman HM (1986) A cyclic AMP- and phorbol ester-inducible DNA element. Nature 323: 353–6

Croll NA, Smith JM, Zuckerman BM (1977) The aging process of the nematode Caenorhabditis elegans in bacterial and axenic culture. Exp Aging Res 3: 175–89

da Graca LS, Zimmerman KK, Mitchell MC, Kozhan-Gorodetska M, Sekiewicz K, Morales Y, Patterson GI (2004) DAF-5 is a Ski oncoprotein homolog that functions in a neuronal TGF beta pathway to regulate C. elegans dauer development. Development 131: 435–46

Fusco G, Minelli A (2010) Phenotypic plasticity in development and evolution: facts and concepts. Introduction. Philos Trans R Soc Lond B Biol Sci 365: 547–56

Golden JW, Riddle DL (1984) The Caenorhabditis elegans dauer larva: developmental effects of pheromone, food, and temperature. Dev Biol 102: 368–78

Jeong PY, Jung M, Yim YH, Kim H, Park M, Hong E, Lee W, Kim YH, Kim K, Paik YK (2005) Chemical structure and biological activity of the Caenorhabditis elegans dauer-inducing pheromone. Nature 433: 541–5

Kaplan F, Srinivasan J, Mahanti P, Ajredini R, Durak O, Nimalendran R, Sternberg PW, Teal PE, Schroeder FC, Edison AS, Alborn HT (2011) Ascaroside expression in Caenorhabditis elegans is strongly dependent on diet and developmental stage. PLoS One 6: e17804

Kim DW, Kempf H, Chen RE, Lassar AB (2003) Characterization of Nkx3.2 DNA binding specificity and its requirement for somitic chondrogenesis. J Biol Chem 278: 27532–9

Kim K, Li C (2004) Expression and regulation of an FMRFamide-related neuropeptide gene family in Caenorhabditis elegans. J Comp Neurol 475: 540–50

Kim K, Sato K, Shibuya M, Zeiger DM, Butcher RA, Ragains JR, Clardy J, Touhara K, Sengupta P (2009) Two chemoreceptors mediate developmental effects of dauer pheromone in C. elegans. Science 326: 994–8

Kim SJ, Jeang KT, Glick AB, Sporn MB, Roberts AB (1989) Promoter sequences of the human transforming growth factor-beta 1 gene responsive to transforming growth factor-beta 1 autoinduction. J Biol Chem 264: 7041–5

Kimura KD, Tissenbaum HA, Liu Y, Ruvkun G (1997) daf-2, an insulin receptor-like gene that regulates longevity and diapause in Caenorhabditis elegans. Science 277: 942–6

Kimura Y, Corcoran EE, Eto K, Gengyo-Ando K, Muramatsu MA, Kobayashi R, Freedman JH, Mitani S, Hagiwara M, Means AR, Tokumitsu H (2002) A CaMK cascade activates CRE-mediated transcription in neurons of Caenorhabditis elegans. EMBO Rep 3: 962–6

Lee JS, Shih PY, Schaedel ON, Quintero-Cadena P, Rogers AK, Sternberg PW (2017) FMRFamide-like peptides expand the behavioral repertoire of a densely connected nervous system. Proc Natl Acad Sci U S A 114: E10726–E10735

Li C, Kim K (2014) Family of FLP Peptides in Caenorhabditis elegans and Related Nematodes. Front Endocrinol (Lausanne) 5: 150

Massague J (2008) TGFbeta in Cancer. Cell 134: 215–30

McGrath PT, Xu Y, Ailion M, Garrison JL, Butcher RA, Bargmann CI (2011) Parallel evolution of domesticated Caenorhabditis species targets pheromone receptor genes. Nature 477: 321–5

Meisel JD, Panda O, Mahanti P, Schroeder FC, Kim DH (2014) Chemosensation of bacterial secondary metabolites modulates neuroendocrine signaling and behavior of C. elegans. Cell 159: 267–80

Neal SJ, Kim K, Sengupta P (2013) Quantitative assessment of pheromone-induced Dauer formation in Caenorhabditis elegans. Methods Mol Biol 1068: 273–83

Nolan KM, Sarafi-Reinach TR, Horne JG, Saffer AM, Sengupta P (2002) The DAF-7 TGF-beta signaling pathway regulates chemosensory receptor gene expression in C. elegans. Genes Dev 16: 3061–73

Park J, Choi W, Dar AR, Butcher RA, Kim K (2019) Neuropeptide Signaling Regulates Pheromone-Mediated Gene Expression of a Chemoreceptor Gene in C. elegans. Mol Cells 42: 28–35

Patterson GI, Koweek A, Wong A, Liu Y, Ruvkun G (1997) The DAF-3 Smad protein antagonizes TGF-beta-related receptor signaling in the Caenorhabditis elegans dauer pathway. Genes Dev 11: 2679–90

Projecto-Garcia J, Biddle JF, Ragsdale EJ (2017) Decoding the architecture and origins of mechanisms for developmental polyphenism. Curr Opin Genet Dev 47: 1–8

Ren P, Lim CS, Johnsen R, Albert PS, Pilgrim D, Riddle DL (1996) Control of C. elegans larval development by neuronal expression of a TGF-beta homolog. Science 274: 1389–91

Savage-Dunn C, Padgett RW (2017) The TGF-beta Family in Caenorhabditis elegans. Cold Spring Harb Perspect Biol 9

Schackwitz WS, Inoue T, Thomas JH (1996) Chemosensory neurons function in parallel to mediate a pheromone response in C. elegans. Neuron 17: 719–28

Srinivasan J, Kaplan F, Ajredini R, Zachariah C, Alborn HT, Teal PE, Malik RU, Edison AS, Sternberg PW, Schroeder FC (2008) A blend of small molecules regulates both mating and development in Caenorhabditis elegans. Nature 454: 1115–8

Vellichirammal NN, Gupta P, Hall TA, Brisson JA (2017) Ecdysone signaling underlies the pea aphid transgenerational wing polyphenism. Proc Natl Acad Sci U S A 114: 1419–1423

von Reuss SH, Bose N, Srinivasan J, Yim JJ, Judkins JC, Sternberg PW, Schroeder FC (2012) Comparative metabolomics reveals biogenesis of ascarosides, a modular library of small-molecule signals in C. elegans. J Am Chem Soc 134: 1817–24

Wang J, Kim SK (2003) Global analysis of dauer gene expression in Caenorhabditis elegans. Development 130: 1621–34

